# Computational Pan-genome Mapping and pairwise SNP-distance improve Detection of *Mycobacterium tuberculosis* Transmission Clusters

**DOI:** 10.1101/752782

**Authors:** Christine Jandrasits, Stefan Kröger, Walter Haas, Bernhard Y. Renard

**Affiliations:** Bioinformatics Unit, Robert Koch Institute, Berlin, Germany; Respiratory Infections Unit, Robert Koch Institute, Berlin, Germany

## Abstract

Next-generation sequencing based base-by-base distance measures have become an integral complement to epidemiological investigation of infectious disease outbreaks. This study introduces PANPASCO, a computational pan-genome mapping based, pairwise distance method that is highly sensitive to differences between cases, even when located in regions of lineage specific reference genomes. We show that our approach is superior to previously published methods in several datasets and across different *Mycobacterium tuberculosis* lineages, as its characteristics allow the comparison of a high number of diverse samples in one analysis - a scenario that becomes more and more likely with the increased usage of whole-genome sequencing in transmission surveillance.

**Author summary:** Tuberculosis still is a threat to global health. It is essential to detect and interrupt transmissions to stop the spread of this infectious disease. With the rising use of next-generation sequencing methods, its application in the surveillance of *Mycobacterium tuberculosis* has become increasingly important in the last years. The main goal of molecular surveillance is the identification of patient-patient transmission and cluster detection. The mutation rate of *M. tuberculosis* is very low and stable. Therefore, many existing methods for comparative analysis of isolates provide inadequate results since their resolution is too limited. There is a need for a method that takes every detectable difference into account. We developed PANPASCO, a novel approach for comparing pairs of isolates using all genomic information available for each pair. We combine improved SNP-distance calculation with the use of a pan-genome incorporating more than 100 *M. tuberculosis* reference genomes for read mapping prior to variant detection. We thereby enable the collective analysis and comparison of similar and diverse isolates associated with different *M. tuberculosis* strains.

## Introduction

Genotyping and sequencing methods have revolutionized infectious disease surveillance. So called molecular surveillance - molecular data in combination with classical epidemiological data - allows the investigation of the transmission of disease within the population and the sensitive detection of outbreaks. The employed methods shifted from fingerprinting, e.g. variable number tandem repeat (VNTR) methods, and sequence-based genotyping assays, such as bacterial multilocus sequence typing (MLST) to next-generation sequencing (NGS) based whole genome sequencing (WGS) in the recent years [1]. WGS gives access to all genetic information and enables studies on phylogeny, geographical spread of lineages, strain-specific differences, virulence and drug resistance [2,3]. WGS allows for the comparison of pathogens on the level of single nucleotide polymorphisms (SNP) and thus is particularly useful for the transmission analysis of stable genomes with low mutation rates such as *Mycobacterium tuberculosis*.

Tuberculosis (TB) is one of the oldest communicable diseases in humankind and can be dated back until 8000 BCE [4]. Although Robert Koch’s characterization of the bacteria is known for more than hundred years, no effective way to eliminate TB has been found and *M. tuberculosis* is still one of the deadliest pathogens worldwide. Between 2000 and 2016 TB caused 53 million deaths. The WHO estimates more than 1.7 million deaths (including HIV coinfection) and 10.4 million new TB infections in 2016 only [5]. Since vaccination is unavailable, only breaking transmission chains and successful treatment can prevent new infections and decrease the spread to finally eliminate TB, which it is the global goal until 2050 [6]. Here, multi-resistant (MDR-) TB is of special interest with about half a million new cases in 2017. MDR-TB alone is responsible for one third of all antimicrobial resistance deaths wordwide [7] and part of WHO’s Ten threats to global health in 2019 list in the context of antimicrobial resistance [8].

In recent outbreak investigations WGS was an indispensable tool of outbreak detection and transmission analysis and is replacing genotyping (mycobacterial interspersed repetitive unit - variable number tandem repeat, MIRU-VNTR) as method of choice in low incident countries like most West-European countries [9–11]. WGS can validate participation of individuals to a common transmission event and thus associate TB cases that do not have a clear epidemiological link or vice versa [9,12,13]. With its high resolution WGS outperforms other molecular typing methods as MIRU-VNTR, Spoligotyping or RFLP [14]. Furthermore, NGS technology granted access to phylogeny, geographical distribution of different *M. tuberculosis* strains and the dynamics of *M. tuberculosis* evolution [12]. Strain differentiation was of specific interest for the last decade and several studies showed how WGS outperforms other genotyping methods for detecting recent transmissions [15,16] and clusters [17,18]. WGS enables base-by-base comparison between two samples with exact distinction of identical and non-identical regions. Reference genomes like the laboratory strain H37Rv allow for comparison of multiple samples by comparing their sequences to these. Over the years a variety of additional reference genomes based on clinical strains with specific drug resistance patterns or specific to certain geographic appearance were created. Thereby, individual reference genomes help to match samples or sample groups like outbreak clusters [2,19,20].

Various methods for measuring the distance between samples using SNPs obtained with NGS are proposed. Different strategies for making distance analysis less complex like core-genome MLST [21] or whole genome MLST [22] have been described. However, these strategies use a gene-by-gene comparison approach. As the molecular clock of *M. tuberculosis* is considered as very low with 0.3-0.5 mutations per genome per year [23,24], the MLST approaches can be insufficient to investigate transmission patterns in clusters or reconstruct direct transmission links. For this reason, we, as well as many outbreak analyses [9–11,16,17,24–27], focus on SNP-counting methods as they retain the highest resolution for patient-patient transmission. We assessed ten exemplary studies on detection of *M. tuberculosis* transmission clusters [10,11,16,17,24–29]. In eight of them the *M. tuberculosis* strain H37Rv (NC_000962.3) is used as the reference genome to identify sample-specific variations. In order to achieve this, samples of patients’ bacteria are sequenced and the resulting reads are mapped to the reference genome with commonly used short read aligners. Then, variant calling tools are used to detect single nucleotide polymorphisms in the sample data. For the majority of studies the main steps after variant calling are similar: SNPs are filtered using high quality standards such as a minimum number of reads mapped to the variant site (5-10 reads), with a high percentage of these reads supporting the variant (75%). In most of these studies additional regions of low confidence, such as repetitive regions, known resistance mutations, regions with more than one SNP, insertion or deletion within 12 bp of the SNP are identified and variant calls within these regions are excluded.

However, the described methods vary in how uncovered sites or low-quality variant calls are handled - they are either completely excluded from the analysis and all comparisons (e.g. [10, 11]) or the low-quality base calls and uncovered regions are ignored and substituted with the reference sequence [17]. Some studies used pairwise comparisons for a small set of samples [28,29]. Several of these strategies are facilitated by the BugMat software [30].

Considering these studies, the questions that are posed for standardizing WGS-based molecular surveillance analyses are: Which genome should the reads be aligned to? How can regions with missing or low-quality information be handled? In alignment-based whole genome sequencing analyses, the choice of the reference genome predetermines the results of subsequent analysis [31]. There is a need to avoid this bias and include multiple reference sequences into the analysis e.g. by using a pan-genome [32]. In this context the term pan-genome describes a set of associated (whole genome) sequences rather than the core and dispensable genes of all strains of an organism. The method of comparison of samples should consider all high-quality variants while excluding regions of low quality for each sample.

We present a new approach, PANPASCO (PAN-genome based PAirwise SNP COmparison), that combines an improved pairwise distance measure, that allows the comparison and clustering of a large number of diverse samples with the use of a computational pan-genome reference sequence. PANPASCO considers each detectable difference between pairs of samples, without sacrificing the ability to resolve intra-cluster patient-patient relations.

## Results

We developed PANPASCO, a novel method to determine the distance between samples based on SNP differences. We compare samples in a pairwise manner, considering all variant sites of high-quality for each pair. These high-quality sites are identified using an NGS variant calling workflow and a five step variant quality filter. To enable the comparison of pairs of samples with differing amount of missing data, the number of low-quality sites and regions with missing information is also incorporated into the distance measure in a normalization step. To minimize the information loss and avoid the problem of identifying each best fitting reference genome per sample, PANPASCO uses a computational pan-genome built from 146 *M. tuberculosis* genomes with seq-seq-pan [33]. This computational pan-genome is about 18% (<1 Mbp) longer than a single *M. tuberculosis* genome, e.g. the H37Rv strain, and contains all genomic regions shared by and specific to each included genome. We use the computational pan-genome sequence in place of a lineage specific reference genome in our mapping and variant calling workflow. When comparing samples using their SNP difference, we can therefore also include SNPs that occur in regions that are not part of commonly used reference strains (for details see Methods).

Several studies investigated the mutation rate for *M. tuberculosis* and evaluated the SNP-based difference cutoffs for detecting transmission between patients and within clusters ranging from 3 to 14 SNPs [10,24–26,34]. Here, we chose a conservative definition that assigns samples with a distance of less than 13 SNPs into transmission clusters and thereby distinguishes them from unrelated samples [10].

We compared PANPASCO to two other commonly used strategies for detecting SNP-based difference, where variant sites and regions with missing information in one of the samples are either completely excluded from the analysis [10,11] or are ignored and therefore substituted with the reference sequence when samples are compared [17] (referred to as exclusion method and substitution method). For these two methods we use the *M. tuberculosis* H37Rv strain as reference genome as described in the respective publications [10,11,17]. Fig 1 shows how these methods work in detail and their individual characteristics: SNP differences in pairs of samples are missed if they are located in regions with missing information in unrelated samples with the exclusion method, while artificial differences are introduced by substituting missing data with the reference sequence when using the substitution method. Fig 1 also shows how all differences between pairs of samples are detected and incorporated in the distance measure with PANPASCO.

**Fig 1.**
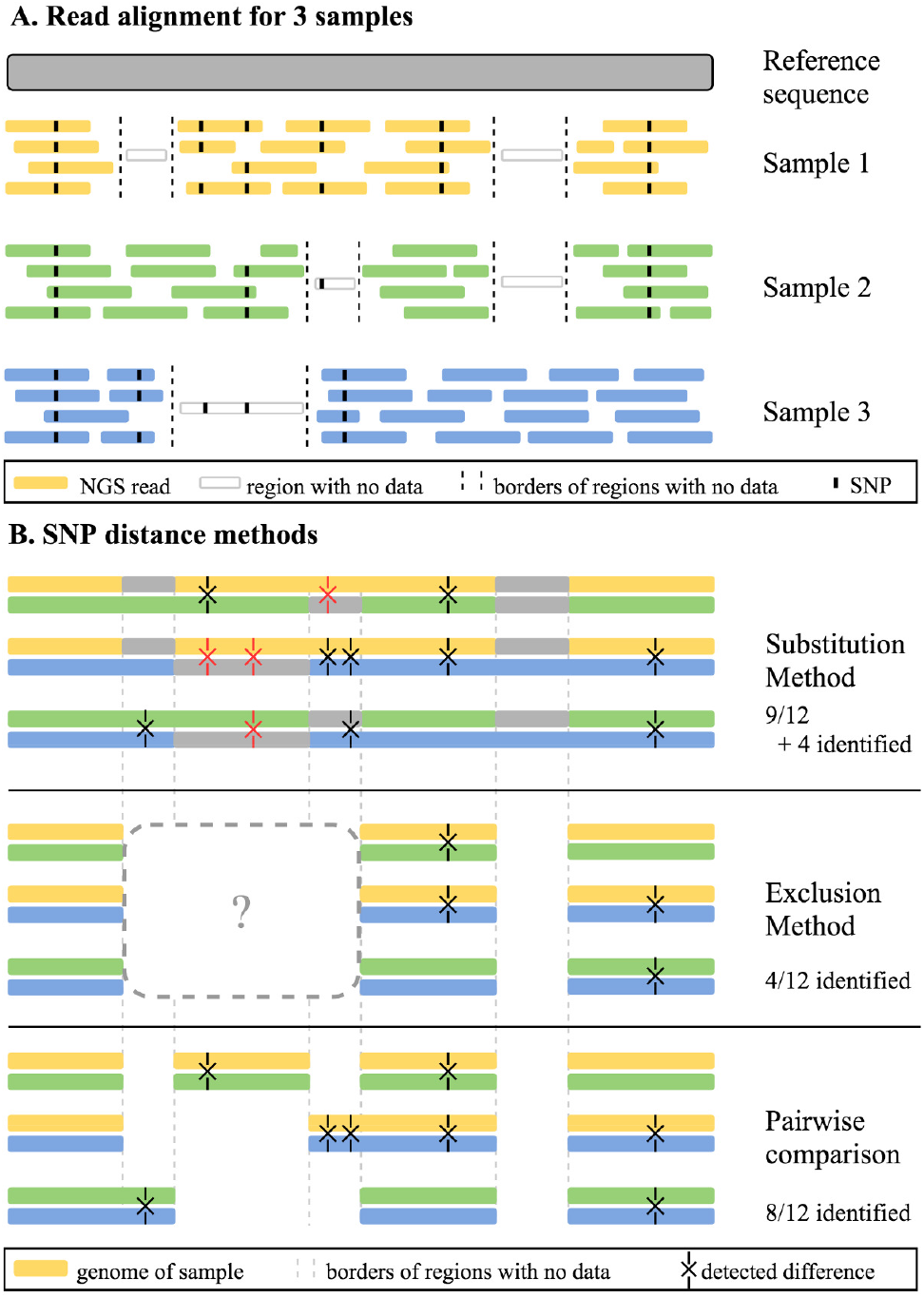
Description and comparison of three SNP distance methods. (A) Schematic representation of the next-generation sequencing (NGS) reads of three samples aligned to a reference sequence. Reference sequence is depicted in gray, while samples are colored yellow, green and blue. Dashed lines represent borders of regions of the samples that are not covered by any NGS reads. Transparent reads represent the true genome of the samples in uncovered regions. Reads reveal single nucleotide polymorphisms (SNPs) when comparing the samples to the reference sequence (depicted as black points on the reads). SNPs also occur in uncovered sequences of samples, these can not be detected using the aligned reads. In sum, the samples have 12 differences, while only 8 can be identified with the available reads. (B) Representation of compared sequence parts using three different distance measuring methods. Each methods compares pairs of samples. Samples are depicted as whole genomes in yellow, blue, and green. Differences between samples are marked with X, true differences are colored black and incorrect ones are colored red. With the substitution method, missing parts in the samples are replaced with the corresponding parts of the reference sequence (in gray). This leads to incorrectly identified differences, where the true, but uncovered sequence of the samples differs from the reference sequence. Using the exclusion method, all uncovered regions of all samples are excluded in all comparisons, e.g. an uncovered region in Sample 1 influences the comparison of Samples 2 and 3. This leads to overlooking differences between samples, in this example only 4 of 12 detectable differences were identified. The pairwise SNP comparison method used in PANPASCO determines all comparable regions for each pair of samples. Uncovered regions of each pair are excluded for pairwise comparison. This way all detectable differences are identified, without introducing additional, incorrect ones.

### Experimental setup

To evaluate the different SNP-counting strategies, we compare the number of transmission cluster links identified by each method in three different datasets. For this, we classify all links between samples into transmission cluster links (SNP difference of less than 13) and unrelated links following previously published results [10] and group samples into transmission clusters. To be assigned to a transmission cluster the distance of one sample to only one of the other samples has to be classified as transmission cluster link, therefore unrelated samples can belong to the same transmission cluster when they are connected by other samples.

We created a simulation dataset with the properties (e.g. number of mutations and coverage distribution) of real datasets [10,35,36] (see S1 Appendix). It includes 20 transmission clusters with 3 to 55 samples per cluster. Samples were simulated from four different genomes including the commonly used *M. tuberculosis* reference strain H37Rv, with five clusters for each genome (for details see Supplementary Methods in S1 Appendix and S1 Table). With this simulation we compare the accuracy of the classification of sample links of the different methods.

We also evaluated PANPASCO’s performance for the analysis of two published datasets, to demonstrate the relevance of our improved distance measure. We chose a study with a large national dataset of 217 samples, with one transmission cluster described in detail [10] and a dataset with a small number of patients [11] (referred to as UKTB (United-Kingdom-TB) and RAGTB (Romania-Austria-Germany-TB), respectively).

### Simulation dataset results

Applying the exclusion method to the simulation dataset results in a high number of predicted transmission cluster links. The sensitivity is very high (1.000), as there are no false negative classifications, however the specificity (0.782) and F1-Score (0.326) are low. In detail, the strategy of excluding low-quality variant sites found in any sample from the transmission analysis results in a high number of false positive classifications - a transmission is assumed where there is no relation between the samples (Table 1).

**Table 1.**
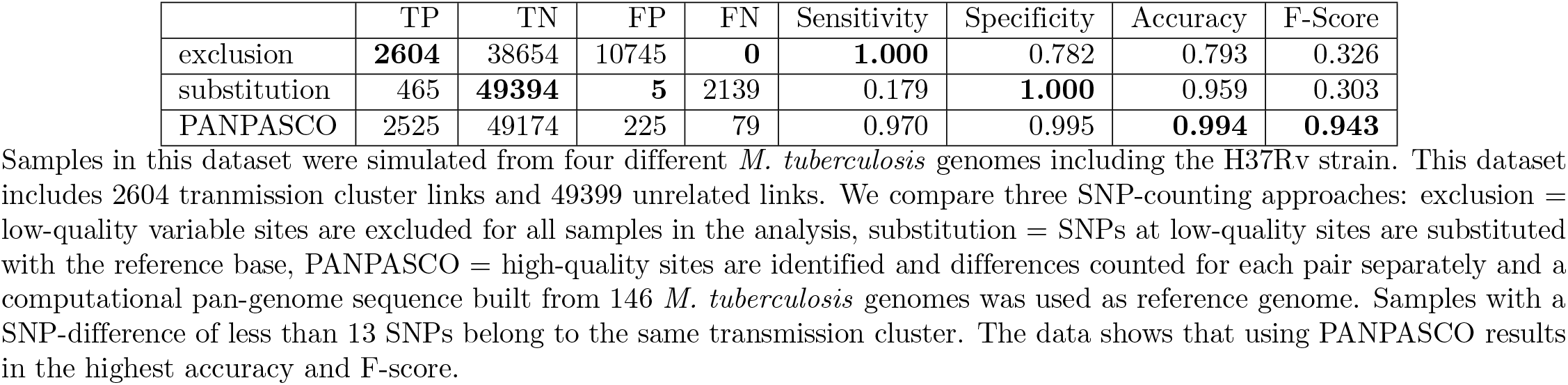
Comparison of SNP-counting methods in all clusters of the simulation dataset.

|  | TP | TN | FP | FN | Sensitivity | Specificity | Accuracy | F-Score |
| --- | --- | --- | --- | --- | --- | --- | --- | --- |
| exclusion | **2604** | 38654 | 10745 | **0** | **1.000** | 0.782 | 0.793 | 0.326 |
| substitution | 465 | **49394** | **5** | 2139 | 0.179 | **1.000** | 0.959 | 0.303 |
| PANPASCO | 2525 | 49174 | 225 | 79 | 0.970 | 0.995 | **0.994** | **0.943** |

Samples in this dataset were simulated from four different *M. tuberculosis* genomes including the H37Rv strain. This dataset includes 2604 transmission cluster links and 49399 unrelated links. We compare three SNP-counting approaches: exclusion = low-quality variable sites are excluded for all samples in the analysis, substitution = SNPs at low-quality sites are substituted with the reference base, PANPASCO = high-quality sites are identified and differences counted for each pair separately and a computational pan-genome sequence built from 146 *M. tuberculosis* genomes was used as reference genome. Samples with a SNP-difference of less than 13 SNPs belong to the same transmission cluster. The data shows that using PANPASCO results in the highest accuracy and F-score.

The substitution method classifies a high number of links between samples as unrelated. This results in many true negative classifications (specificity = 1.000), but also in many false negative ones (sensitivity = 0.179). Substituting regions of low quality with the sequence of the reference genome introduces artificial differences between the samples and therefore this method cannot be used to identify transmission links anymore, as the relation between samples is obscured (Table 1).

Transmission cluster links can be accurately identified using PANPASCO resulting in the highest accuracy and F-Score for the simulation dataset (>0.99 and >0.94, respectively, Table 1).

To analyze the influence of the reference genome on the results of the different methods, we group all samples by the genome used for simulation and compare the results. The 74 samples of the first set of clusters (C1-C5) were simulated from the *M. tuberculosis* H37Rv strain, which is commonly used for SNP distance analyses [10,11,17]. The samples in the rest of the clusters are simulated from three other genomes that are increasingly different from the H37Rv strain (see S2 Table) and therefore increase the diversity between analyzed samples. For details on all genomes used for simulation see Supplementary Methods in S1 Appendix and S1 Table.

Due to the concept of the exclusion method, results change with the number of samples and the sequence diversity among the samples that are included in the analysis. For this reason we used the exclusion method in two ways, once including only the clusters simulated from the respective genome and once including all samples but show the results for the respective samples only. The differing results for these strategies are clearly evident in Table 2. Using the exclusion method for analyzing groups of very similar samples works well. In contrast, when more samples, originating from different genomes are included, more genomic regions are excluded from the analysis and therefore less differences are detected. This results in a strong increase of false-positive classifications of sample links and therefore a large decrease in specificity and accuracy.

**Table 2.** Comparison of SNP-counting methods for simulated samples grouped by the genome used for simulation.

| Genome | Method | TP | TN | FP | FN | Sensitivity | Specificity | Accuracy | F-Score |
| --- | --- | --- | --- | --- | --- | --- | --- | --- | --- |
| H37Rv | exclusion, cluster samples* | 499 | 1915 | 287 | 0 | **1.000** | 0.870 | 0.894 | 0.777 |
|  | exclusion* | 499 | 0 | 2202 | 0 | **1.000** | 0.000 | 0.185 | 0.312 |
|  | substitution | 463 | 2197 | 5 | 36 | 0.928 | **0.998** | 0.985 | 0.958 |
|  | PANPASCO | 489 | 2187 | 15 | 10 | 0.980 | 0.993 | **0.991** | **0.975** |
| MDRMA208 | exclusion, cluster samples* | 464 | 2279 | 260 | 0 | **1.000** | 0.898 | 0.913 | 0.781 |
|  | exclusion* | 464 | 0 | 2539 | 0 | **1.000** | 0.000 | 0.155 | 0.268 |
|  | substitution | 2 | 2539 | 0 | 462 | 0.004 | **1.000** | 0.846 | 0.009 |
|  | PANPASCO | 425 | 2520 | 19 | 39 | 0.916 | 0.993 | **0.981** | **0.936** |
| HKBS1 | exclusion, cluster samples* | 440 | 1531 | 109 | 0 | **1.000** | 0.934 | 0.948 | 0.890 |
|  | exclusion* | 440 | 0 | 1640 | 0 | **1.000** | 0.000 | 0.212 | 0.349 |
|  | substitution | 0 | 1640 | 0 | 440 | 0.000 | **1.000** | 0.788 | 0.000 |
|  | PANPASCO | 425 | 1616 | 24 | 15 | 0.966 | 0.985 | **0.981** | **0.956** |
| TB282 | exclusion, cluster samples* | 1201 | 3679 | 685 | 0 | **1.000** | 0.843 | 0.877 | 0.778 |
|  | exclusion* | 1201 | 0 | 4364 | 0 | **1.000** | 0.000 | 0.216 | 0.355 |
|  | substitution | 0 | 4364 | 0 | 1201 | 0.000 | **1.000** | 0.784 | 0.000 |
|  | PANPASCO | 1186 | 4197 | 167 | 15 | 0.988 | 0.962 | **0.967** | **0.929** |

Comparison of sample link classification for all samples of the simulated dataset grouped by the genome used for simulation shows the impact of reference genome for the different methods. For all samples the same reference genome was used in the analysis: the *M. tuberculosis* H37Rv strain for the exclusion and substitution method and the computational pan-genome reference sequence for PANPASCO. The results show a great reduction in accuracy and F-Score for the groups of samples that were simulated from genomes other than the H37Rv strain for the substitution method and the exclusion method when using all samples in the distance calculation. PANPASCO achieves very good classifications with no difference between the groups of samples.
\* Results change with the number and diversity of samples when using the exclusion method. We show the results for each group, including only the respective cluster samples or all simulated samples in the analysis.

The choice of reference genome strongly affects the results obtained with the substitution method. Substituting regions of low quality with the reference genome works well only if the analyzed samples are mapped to the best fitting reference genome: almost all transmission cluster links between samples simulated from genomes other than the H37Rv strain were misclassified as unrelated (Table 2).

Using PANPASCO, the classification results do not differ between the four groups of samples. We achieved the highest accuracy and F-Score (>0.96 and >0.92, respectively) for all links between samples (Table 2). This detailed analysis shows the advantage of our approach: links between all samples in a large, diverse dataset are classified equally well, independent of the number of included samples or strains, as all SNPs, even those in strain specific genomic regions are taken into account for measuring the distance between pairs of samples.

Detailed inspection of sample links for each simulated cluster (S4 Table) confirms the shortcomings of the other methods described in Fig 1: Only inter-cluster links were correctly classified as unrelated by the exclusion method, but all samples within clusters were reported as closely related, even if the clusters contained samples with no relation. For example, in Cluster C1 there are 17 closely related samples (less then 5 SNPs difference), 110 samples with 6-12 SNPs difference and 44 unrelated links and the substitution method reported 154 closely related (S4 Table). This method also reported only four instead of 20 transmission clusters - all samples simulated from the same genome are clustered together. Comparing the number of SNPs detected for each link within each simulated cluster underlines the properties of the other methods (see Fig 2): Due to the diversity of the genomes used for the simulation and the resulting exclusion of genomic regions, most samples within clusters have a reported difference of 0 or 1 with the exclusion method. The opposite is true for the substitution method as most transmission cluster links were reported as being unrelated links (> 12 SNPs).

**Fig 2.**
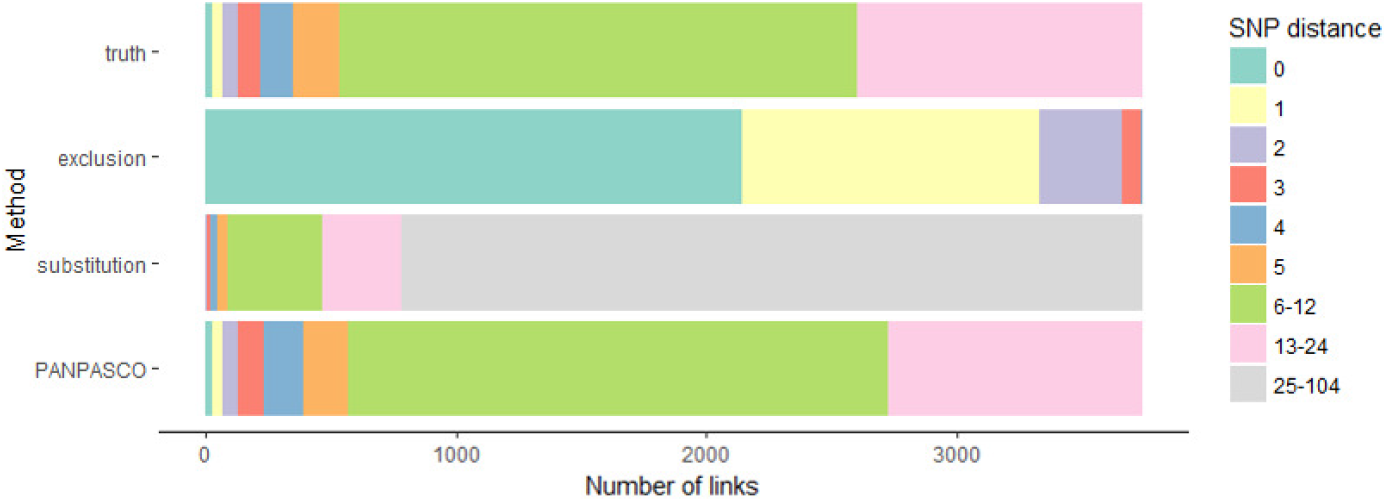
Counting difference between samples within clusters. We select each cluster individually and count the differences between samples within the clusters. Each cluster can contain closely related samples (0 SNPs), transmission cluster links (less than 13 SNPs) and unrelated samples. Samples are assigned to a cluster if the difference is less than 13 SNPs compared to at least one sample within the cluster. We evaluate the frequency of each distance value for the three methods and compare them to the true distance. The exclusion method reports distances of 0-3 SNPs for all samples and with the substitution method all distances are greater than 12. The distribution of distances using PANPASCO closely resemble the true distances in the dataset.

For a complete assessment of the different methods we also used our computational pan-genome reference sequence with the exclusion and substitution methods and PANPASCO with the *M. tuberculosis* H37Rv strain for the analysis of the complete simulation dataset. S5 Table shows that the classification results do not change for the exclusion method. The combination of the substitution method and the pan-genome achieves very poor results as not a single transmission cluster link was detected. PANPASCO works best with the computational pan-genome reference sequence, but also achieves better results than the other two methods using the standard reference genome.

### Real datasets results

Both previously published datasets were analyzed using the computational pan-genome as reference sequence and the mapping and variant calling workflow described in the Supplementary Methods in S1 Appendix.

For the UKTB dataset we focused our comparison on the cluster for which an epidemiological network and nucleotide variants were provided (cluster seven, UKTB7) [10]. This cluster was initially defined by the shared MIRU-VNTR profile of the samples and includes 17 sequenced isolates of ten patients with one central, treatment non-compliant individual. When calculating the SNP distance using PANPASCO, we identified one pair of samples with 4 differing SNPs in addition to the ones described in the original manuscript: P066 and P175. These patients were reported with a distance of 0, indicating a close transmission event or even direct transmission. The minimum spanning tree we calculated with Cytoscape App Spanning Tree [37] based on our SNP differences better matches the described epidemiological network (Fig 3). In contrast to the findings in [10] it is more likely that P076 infected the other two cases, rather than one of these the other one. We investigated the variant sites that we identified and compared them to the published ones. We identified 44 variant sites including all 20 sites listed in [10] (depicted in S6 Table). In the original manuscript the exclusion method was used and the additional 24 sites we identified were excluded because they are not covered in samples assigned to MIRU-VNTR cluster six (data not shown). This lack of coverage is likely explained by the differing phylotypes of cluster six and seven (see Table in [10]).

**Fig 3.**
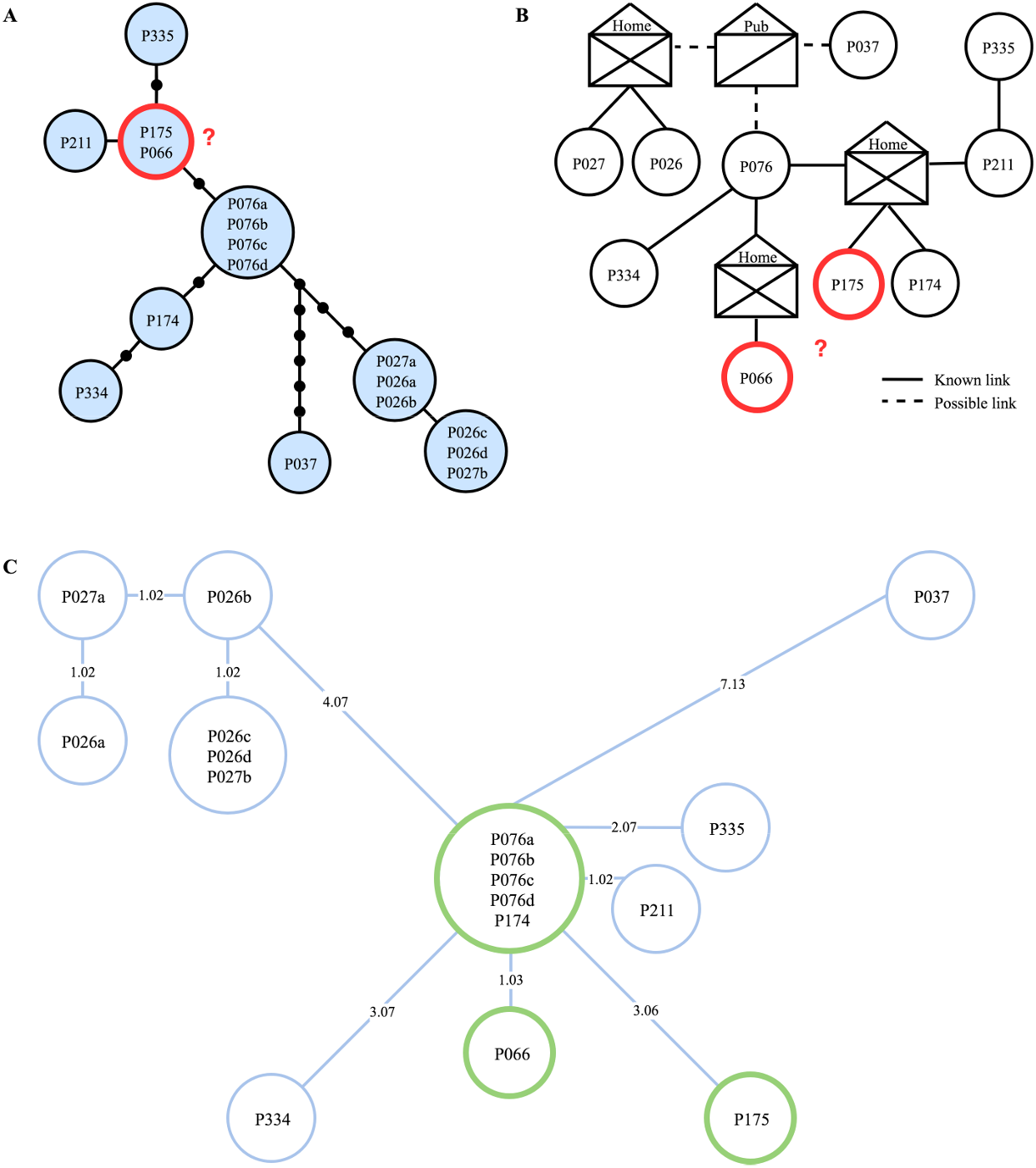
Minimum spanning tree computed with PANPASCO between samples of the UKTB7 dataset. (A) Genetic distances estimated by Walker et al. SNP distances are represented by dots on edges, isolates within blue circles are separated by 0 SNPs. (B) Epidemiological network as published by Walker et al. (C) Minimum spanning tree from distances calculated with PANPASCO. Adjoining isolates are separated by 0 SNPs, edges are labelled with pairwise distance. Transmission links between P066 and P175 are marked in all three parts - more SNPs were detected using PANPASCO and the resulting tree better fits the epidemiological network. ((A) and (B) taken from [10] and edited.)

The second dataset, RAGTB [11] consists of 13 patients and two replicates for one of the patients. We again calculated a minimum spanning tree using distances calculated with PANPASCO (see Fig 4). We identified the same transmission clusters as described in the original manuscript with one exception: We identified more than 12 differing SNPs for patients VI and III. The patients were previously analyzed using the exclusion method, with additional filtering of SNPs associated with drug resistance or located in repetitive regions of the genome [11]. As this is a small dataset, there is not a large difference between the results of our approach and the exclusion method.

**Fig 4.**
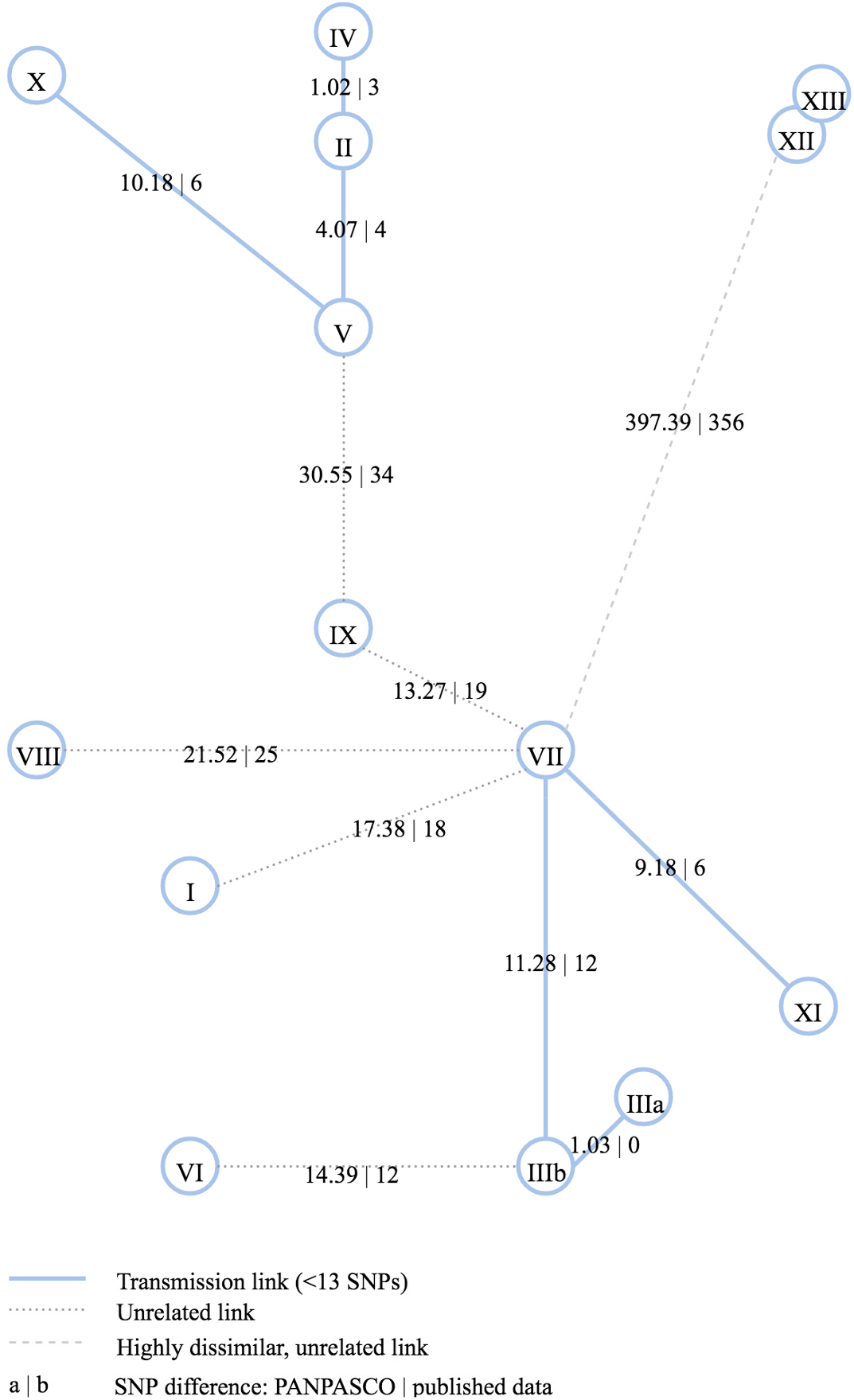
Minimum spanning tree computed from pairwise distances between samples of the RAGTB dataset. Edges are labeled with SNP distances - PANPASCO | published data. The exclusion method and the *M. tuberculosis* H37Rv strain were used in the original publication [11]. Blue lines mark distances of less than 13 SNPs detected with PANPASCO, clustering patients into three transmission clusters and four independent samples. Samples XII and XIII show no difference with both analyses. The comparison of these two distances shows that there is no large difference in the results as this is a small dataset with very similar samples.

## Discussion

We present a new approach for measuring genomic distance of *M. tuberculosis* isolates, integrating commonly used mapping and variant calling methods. Reads of isolates are mapped to a computational pan-genome built from over a hundred *M. tuberculosis* genomes to include strain specific genomic regions in the analysis. SNP differences are evaluated pairwise so all high-quality regions and incorporating regions of low-quality or missing information are considered.

Our method PANPASCO accurately determines transmission clusters within large sets of samples. Previously published and commonly used methods struggle with large, diverse datasets, showing either low specificity or sensitivity when identifying transmissions. Any sort of error is problematic in a disease outbreak investigation. Methods lacking sensitivity result in overlooking transmission and failing outbreak detection. However, also methods lacking specificity produce unsatisfying results. Any health care system can only follow-up on a limited number of possible transmission events and must prioritize its actions and concentrate capacities and efforts. Having large numbers of spurious and incorrect transmission events detected, will automatically impact the availability of resources to focus on the true underlying transmission events within short time.

The major drawback of the exclusion method is its limited resolution, which is driven by the sample with the lowest coverage. Another disadvantage of this method for usage in disease surveillance is that each time a new sample is added to the dataset, the results for previously analyzed isolates change as more regions of the genome have to be excluded to be able to compare the new sample to the dataset. Taking this further, with a rising number of samples at some point, due to random sequencing errors and varying coverage distribution, there will be fewer and fewer sites of the genome with high-quality information available for comparison. This problem is of special interest and high importance regarding long time infectious disease surveillance rather than outbreak investigations and recent transmission analysis.

This problem can be avoided when low-quality or missing regions are substituted with the reference sequence. With the substitution method all detectable difference between samples are considered in the distance calculation. However, many artificial differences are introduced as well. When an *M. tuberculosis* isolate is compared to a reference sequence, usually several hundred SNPs are detected. Within a transmission cluster all but a few of these SNPs are the same. When a subset of these sites cannot be detected due to missing or low-quality reads and are substituted with the reference base, they are counted as differences between the samples and transmissions cannot be detected. The problem increases the more the isolates diverge from the reference sequence.

Strains of *M. tuberculosis* can be assigned to different sub-lineages that differ in many loci and blocks of deletions [36]. Samples of different strains are often analyzed together, choosing one reference genome that naturally represents only one of the lineages adequately [10,16,17,25]. This means, that for a part of the set of samples a suboptimal reference is used for mapping and variant calling. Nevertheless, identifying the best fitting reference sequence for each sample is no optimal solution. Clusters within a dataset and datasets from different studies will not be comparable with each other. The computational pan-genome approach solves this problem as it allows the integration of genomic information of several *M. tuberculosis* strains. The computational pan-genome enables the detection of SNPs in the core genome and in strain specific genomic regions at the same time.

We showed the specific weaknesses of the exclusion and substitution approach by applying them to a simulation dataset that was constructed to resemble a real dataset. We demonstrate the superior classification result of combining the usage of the computational pan-genome reference and the pairwise comparison. Using PANPASCO to analyze previously published datasets provided additional insight for the epidemiological investigation of these transmission clusters.

Having differing numbers of comparable sites for each pair in the analysis, complicates distance calculation. To account for the fact that fewer differences can be detected with less comparable sites we project the number of SNPs found for each pair to the full length of the genome, e.g. detecting 2 differences within 2 Megabases is different from detecting 2 differences in 4 Megabases using a normalized score. For identifying comparable sites and incorporating missing regions into the distance measure we keep track of all, instead of only variable high-quality sites of all samples. In this normalization step we work with the assumption that SNPs in samples are equally distributed over the length of the computational pan-genome. Mutation hot spots and the frequency of variants in the genomes used for the computational pan-genome can be taken into account for a more accurate normalized score.

Today, WGS-based molecular surveillance of MTB is established in a number of low-incident countries, e.g. USA, UK, Netherlands. Such systems allow for event specific adaption of public health action, patient care, medication and treatment based on pathogen specifics like resistance, virulence and actual spread (outbreak size). Early detection of antibiotic resistance and prevention of further transmission are one of the main tasks on the path to TB elimination. This implies a large number of samples that has to be analyzed - e.g. in Germany there were more than 4000 cases (4099 of 5915 reported cases; corresponding to 83.4%) laboratory confirmed by culture of tuberculosis in 2016 [38]. Nowadays, higher mobility and worldwide migration cause a larger geospatial spread of specific pathogens and increase the diversity of samples.

Several studies analyzed and assessed the integration of WGS into routine tuberculosis diagnosis and investigation [10,39–41]. The authors show the added benefit of using SNP-based analysis in transmission cluster and drug resistance detection in large groups of patients. However, while they discuss the importance of common standards for sequencing techniques and quality, the implications of integrating a large number of samples across different lineages in the same analysis are not addressed.

Balancing sensitivity and specificity is key for the analysis of large and diverse groups of samples during outbreak investigations and in TB surveillance, when it is of importance to find each case and expensive to investigate large numbers of false positives. PANPASCO can contribute to achieving these goals by usage of pan-genomic references and improved pairwise SNP-distances.

## Materials and Methods

### *PAN* - Computational Pan-genome Mapping

In mapping based whole genome sequencing analyses, the choice of the reference genome can have significant impact on the results [31]. For this reason we built a computational pan-genome from 146 *M. tuberculosis* genomes available in NCBI RefSeq by February 17th, 2018 with seq-seq-pan [33]. seq-seq-pan aligns all genomes in an iterative way, adding new genomic content step by step. This resulted in a computational pan-genome sequence with 5,205,216 bp (an increase of about 18% compared to 4,411,532 bp of the commonly used *M. tuberculosis* H37Rv strain) and contains all genomic regions shared by and specific to each included genome (see list of genomes in S7 Table). We use this computational pan-genome sequence as reference sequence with a pipeline that includes various tools for quality control, mapping and variant calling and filtering, with bwa mem [42] for read alignment and GATK for variant detection [43] (Supplementary Methods in S1 Appendix and S1 Fig). Scripts for the whole analysis workflow are provided at https://gitlab.com/rki_bioinformatics/panpasco.

### *PASCO* - Pairwise SNP Comparison

The first step of distance calculation is the identification of high-quality SNPs. For this we use several filters to identify uncovered, low-quality and ambiguous sites for all samples (Supplementary Methods in S1 Appendix and S1 Fig). Then, we compare all samples *pairwise*, taking into account the set of all variant sites of high-quality (*S*) in the genomes of a pair of samples (*x*_1_ and *x*_2_) compared to a reference genome. This way we do not lose information about differing bases that are located in low-quality regions of other, unrelated samples. Opposed to the previously published exclusion and substitution methods, we end up with different numbers of sites for each comparison, due to differing number and length of low-quality regions in each sample. To account for this difference we normalize the SNP count by this number of compared sites. This score reflects the SNP difference per base.

We also determine common reference genome sites of the samples (*G*). For this we compare the low-quality regions of the samples with the whole genome alignment (WGA) that forms the computational pan-genome sequence. The common reference genome sites are composed of high-quality sites of the samples and the low-quality sites that do not overlap with gaps in the WGA of the computational pan-genome. Overlaps with gaps in the WGA indicate that the reason for lack of coverage are strain differences rather than low-quality sequencing (Fig 5).

**Fig 5.**
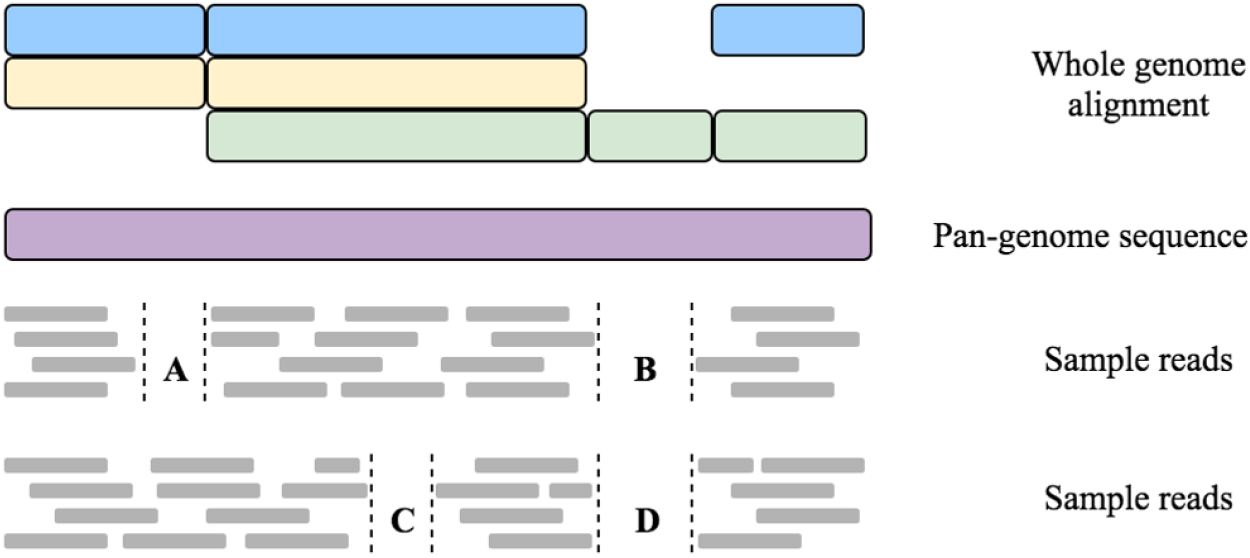
Reads from two samples mapped to a computational pan-genome sequence with regions of zero coverage. Uncovered regions such as A and C are considered to contain as many difference between the samples as found in covered regions. To account for these uncovered regions and the total expected difference between two samples is calculated using the SNP difference per base - derived from regions covered in both samples - and the set of common reference genome sites. This set is composed of all sites of the genome except regions such as B and D. These regions are uncovered in both samples and overlap with gaps in the whole genome alignment (blue, yellow and green) of the strains used to build the computational pan-genome (purple). This indicates that both samples are related to similar strains that both do not contain this specific genomic region, which should therefore not be considered when calculating the expected number of differences for the whole computational pan-genome.

To calculate the expected number of differences for the whole reference genome we multiply the SNP difference per base with the number of common reference genome sites (see Eq (1) and Eq (2)).

We define the distance between the genomes of a pair of samples *x*_1_ and *x*_2_ as

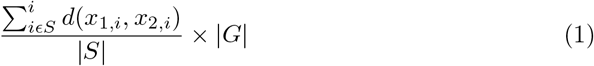

where

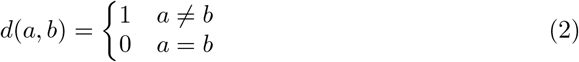

and |*S*| and |*G*| are the number of compared high-quality variant sites and the number of common reference genome sites, respectively.

### Simulation dataset

Following published real datasets (UKTB and RAGTB) we simulated 20 transmission clusters with 3 to 55 samples per cluster, resulting in a total number of 323 samples. We chose four *M. tuberculosis* genomes and assigned five clusters to each of them (S1 Table). For the genomes we chose the *M. tuberculosis* H37Rv strain, commonly used as reference genome, and three increasingly divergent genomes from the set of genomes that built up the computational pan-genome (NC_000962.3, NZ_CP023628.1, NZ_CP002871.1, NZ_CP017920.1). For each cluster we generated intra- and inter-cluster SNPs and we assigned all inter-cluster and a selection of the intra-cluster SNPs to each sample (see S1 Appendix). We estimated the number and length of low coverage regions in the comprehensive UKTB dataset and introduced them into the simulated samples. Short reads were simulated with NEAT [44]. For this purpose we created a variant calling file (VCF) with all assigned SNPs for each sample, including deletions to simulate regions with low coverage (see S1 Appendix for all simulation details).

To create a fair simulated dataset we generated a whole genome alignment of all applied reference genomes with seq-seq-pan [33] to calculate the true distance between the samples (S2 Table). We combined these differences and all simulated SNPs into a distance matrix for all samples, which represents the true distance between all samples. We assessed the number of simulated inter- and intra-cluster SNPs located in regions that are not part of the H37Rv strain by mapping their positions using the coordinate system of the computational pan-genome (S3 Table). Scripts for generation of intra- and intercluster SNPs and deletions are provided at https://gitlab.com/rki_bioinformatics/panpasco.

## Acknowledgments

We would like to thank Pascal Wetzel and Julius Tembrockhaus for assistance in carrying out computational pan-genome related experiments and Lena Fiebig and Andrea Sanchini for providing constructive suggestions and advice on molecular surveillance of tuberculosis.

## Supporting information

**S1 Appendix. Supplementary Methods.** Appendix S1 contains additional details on analyses used in this study. The mapping and variant calling workflow as well as the creation of the simulation dataset are described.

**S1 Fig. Workflow used to analyze the simulation and real datasets.** We divide the task in three parts: read mapping, variant calling and filtering of variants and detection of low-quality regions. We prepare the reads by removing sequence adapters and merging overlapping paired end reads. After that low-quality reads are filtered and reads with ends of low base quality are trimmed. High-quality reads are mapped to a reference and necessary steps for variant calling are taken. We use GATK [43] for variant calling and bedtools [45] to calculate the coverage of the genome. The results are filtered to detect regions of low-quality and high-quality SNPs.

**S1 Table. Description of clusters in the simulation dataset.**

**S2 Table. Description of genomes used for the simulation dataset.** Base differences were counted in a whole-genome alignment of the four genomes. Upper triangular part of table shows hamming distance of sequences in WGA ignoring gaps, while the lower part of the table lists all differences, including gaps.

**S3 Table. Comparison of genomes used in the simulation dataset to the *M. tuberculosis* strain.** We count the simulated SNPs in all simulated samples located on genome specific regions that are not part of the H37Rv genome.

**S4 Table. Details of transmission detection in simulated clusters.** S4 Table provides the counts for transmission cluster links for the exclusion, substitution and pairwise method for mappings to the *M. tuberculosis* H37Rv strain and the computational pan-genome.

**S5 Table. Comparison of SNP-counting methods using different reference genomes.** We used the all three methods (exclusion, substitution, PANPASCO) with the commonly used *M. tuberculosis* H37Rv and the computational pan-genome reference genomes for classification of links between samples in the simulated dataset.

**S6 Table. Comparison of differential SNPs detected in the UKTB7 dataset.** S6 Table contains the full SNP matrix for all samples in the UKTB7 dataset.

**S7 Table. Genomes in computational pan-genome** S7 Table provides detailed description of the genomes used for building the computational *M. tuberculosis* pan-genome including accession numbers and sort order.

