## Supplemental Material for "Computational Pan-genome Mapping and pairwise SNP-distance improve Detection of *Mycobacterium tuberculosis* Transmission Clusters"

#### Read mapping and variant detection workflow

We implemented a workflow for read mapping, variant discovery and detection of low-quality regions for paired-end next generation sequencing samples. It is depicted in S1 Fig.

In this workflow we start with preparing the reads by removing sequence adapters from their ends with Trimmomatic [1] and merging paired-end reads in case they overlap with at least 10 bp with Flash [2]. These steps are necessary in cases where the sequenced DNA fragments are shorter than twice the desired read length. This results in two sets of reads: non-overlapping paired-end reads and merged, longer single reads. Both sets are subjected to quality control with Trimmomatic [1], where reads with bases of low quality are trimmed. In case reads are now shorter than 50 bp they are completely removed from the dataset. Remaining mates of excluded reads are added to the set of single reads.

Each of the sets is mapped to a reference genome with bwa mem [3] and the sets of mapped reads are joined for the following analysis with samtools [4]. Duplicated reads are marked and read groups added with picard tools [5] in preparation for variant detection. Reads with a mapping quality less than 10 are not considered in the following analysis steps.

We use the Genome Analysis Toolkit (GATK, [6]) for variant detection. First we detect confidence scores for all sites, including reference sites with the HaplotypeCaller tool. After that we extract genotypes for each site using the GenotypeGVCFs tool. We call variants with a diploid model to be able to filter mixed base calls by allele frequency. We analyze all sites and separate them in variant sites, positions with uncalled genotypes and high-quality reference sites. Variant sites are then split into single nucleotide polymorphisms (SNPs), small deletions and insertions and structural variants with SelectVariants tool from GATK. We identify SNPs with an allele frequency of at least 75% and where 10 or more reads where used to call the SNP. We also use bedtools [7] to extract regions with less than 10 reads coverage.

By using these filters we separate the data into high-quality SNPs and five sets of low-quality regions with the following criteria:

- positions with less than 10 mapped reads
- positions at deletions
- positions with uncalled genotypes
- SNPs called from less than 10 reads
- SNPs with less than 75% allele frequency

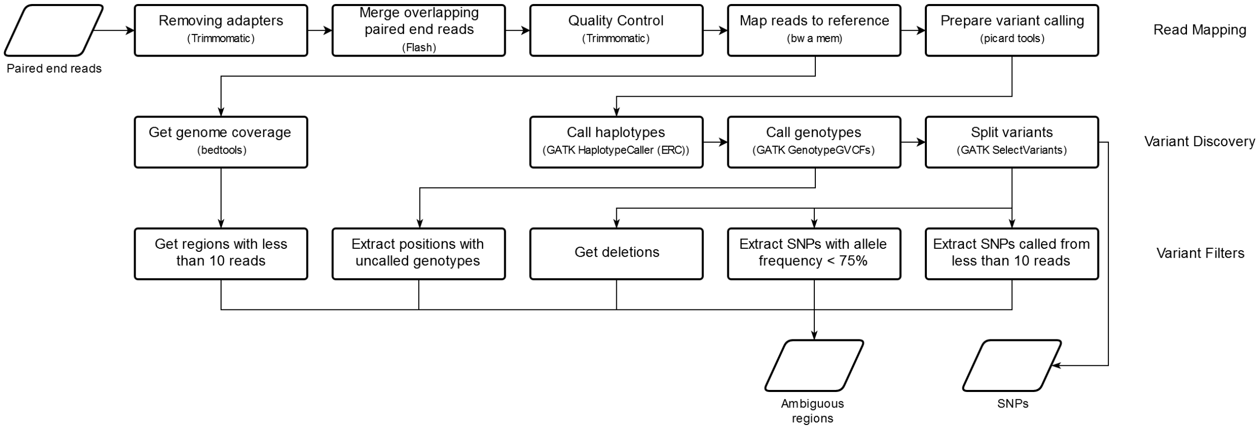


**Fig S1. Workflow used to analyze the simulation and real datasets.** We divide the task in three parts: read mapping, variant calling and filtering of variants and detection of low-quality regions. We prepare the reads by removing sequence adapters and merging overlapping paired end reads. After that low-quality reads are filtered and reads with ends of low base quality are trimmed. High-quality reads are mapped to a reference and necessary steps for variant calling are taken. We use GATK [6] for variant calling and bedtools [7] to calculate the coverage of the genome. The results are filtered to detect regions of low-quality and high-quality SNPs.

### Simulation dataset

The simulation dataset was set up to include several transmission clusters with varying numbers of samples based on different strain genomes, to reflect the properties of a real dataset. We chose the most commonly used *M. tuberculosis* H37Rv strain and three increasingly differing genomes. For the comparison of the genomes we used the whole genome alignment of the 146 *M.tuberculosis* genomes that make up our pan-genome.

We simulated 20 transmission clusters with cluster sizes between 3 and 55. We used a beta distribution in R (v3.3, [8]) with non-negative parameters set to 2 and 9 for a right-skewed distribution of randomly chosen cluster sizes. Number of inter-cluster SNPs were determined for each cluster by random uniform sampling between 6 and 40. A minimum of 6 was chosen so that the distance between two clusters is at least 12 SNPs - the cutoff we chose to separate transmission clusters. The number of intra-cluster SNPs is 1.5 times the cluster size and at least 11 (see cluster sizes and number of inter- and intra-cluster SNPs in S1 Table).

We sampled the positions for all SNPs on the respective genomes between 1 and the genome length without replacement. Then, we assigned inter-cluster SNPs to each sample within the respective cluster. We chose 1 to 11 intra-cluster SNPs for each sample from the set of intra-cluster SNPs assigned to each cluster.

We estimated the number and length of low coverage regions (referred to as *deletions* hereafter) in the comprehensive dataset of [9]. We created a list of all lengths of the detected deletions, including duplicates. For each sample we chose the lengths for 70 to 550 deletions from this list and uniformly sampled their positions on the respective genomes.

This data was used to simulate short reads for each sample with NEAT [10]. For this purpose we created a variant calling file (VCF) with all assigned SNPs and deletions for each sample to simulate mismatches and regions with low coverage. As NEAT cannot handle ambiguous bases within the simulation VCF we replaced them with 'A' in all genomes. As they were only part of deletions this did not influence the simulated short reads. We randomly chose a replacement nucleotide from the set of A, C, T, G for each SNP position.

All read data for the simulated dataset can be downloaded at `https://doi.org/10.5281/zenodo.1346307`.

To account for the differences between the genomes we used for the simulation, we generated a whole genome alignment of the four genomes with seq-seq-pan [11] and counted the number of unequal bases in the alignment (S2 Table). We combined these differences and all simulated SNPs into a distance matrix for all samples, which represents the true distance between all samples.

We assessed the number of simulated inter- and intra-cluster SNPs located in regions that are not part of the H37Rv strain by mapping their positions using the coordinate system of the pan-genome. Several SNPs simulated on the three genomes are not part of the H37Rv strain and therefore can not be detected when using this strain as reference genome (S3 Table).

**Table S1. Comparison of SNP-counting methods in all clusters of the simulation dataset.**

| | | Sensitivity | Specificity | Accuracy | F-Score |
|---|---|---|---|---|---|
| exclusion | H37Rv | **1.000** | 0.782 | 0.793 | 0.326 |
| exclusion | pan-genome | **1.000** | 0.782 | 0.793 | 0.326 |
| substitution | H37Rv | 0.179 | **1.000** | 0.959 | 0.303 |
| substitution | pan-genome | 0.000 | **1.000** | 0.950 | 0.000 |
| PANPASCO | H37Rv | 0.896 | 0.998 | **0.993** | **0.930** |
| PANPASCO | pan-genome | 0.970 | 0.995 | **0.994** | **0.943** |

**Table S2. Description of genomes used for the simulation dataset.**

| | H37Rv | MDRMA2082 | HKBS1 | TB282 |
|---|---|---|---|---|
| H37Rv | 0 | 1046 | 2321 | 2508 |
| MDRMA2082 | 1279 | 0 | 2273 | 2461 |
| HKBS1 | 104466 | 104367 | 0 | 408 |
| TB282 | 132850 | 132766 | 41429 | 0 |

Base differences were counted in a whole-genome alignment of the four genomes. Upper triangular part of table shows hamming distance of sequences in WGA ignoring gaps, while the lower part of the table lists all differences, including gaps.

**Table S3. Comparison of genomes used in the simulation dataset to the *M. tuberculosis* strain.**

| | inter-cluster SNPs | intra-cluster SNPs |
|---|---|---|
| MDRMA2082 | 0 | 1 |
| HKBS1 | 4 | 3 |
| TB282 | 8 | 9 |

We count the simulated SNPs in all simulated samples located on genome specific regions that are not part of the H37Rv genome.

**Table S5. Comparison of SNP-counting methods in all clusters of the simulation dataset.**

| | | Sensitivity | Specificity | Accuracy | F-Score |
|---|---|---|---|---|---|
| exclusion | H37Rv | **1.000** | 0.782 | 0.793 | 0.326 |
| exclusion | pan-genome | **1.000** | 0.782 | 0.793 | 0.326 |
| substitution | H37Rv | 0.179 | **1.000** | 0.959 | 0.303 |
| substitution | pan-genome | 0.000 | **1.000** | 0.950 | 0.000 |
| PANPASCO | H37Rv | 0.896 | 0.998 | **0.993** | **0.930** |
| PANPASCO | pan-genome | 0.970 | 0.995 | **0.994** | **0.943** |

**Table S4, S6 and S7 are provided as separate supplementary files.**
